# Temporal contiguity training does not affect size-tolerant representations in object-selective cortex

**DOI:** 10.1101/791392

**Authors:** Chayenne Van Meel, Hans P. Op de Beeck

**Author notes:** Correspondence: Hans P. Op de Beeck, Laboratory of Biological Psychology, Brain and Cognition, Tiensestraat 102, box 3714, 3000 Leuven, Belgium.

## Abstract

The human visual system has a remarkable ability to reliably identify objects across variations in appearance, such as variations in viewpoint, lighting and size. Here we used fMRI in humans to test whether temporal contiguity training with natural and altered image dynamics can respectively build and break neural size tolerance for objects. Participants (N = 23) were presented with sequences of images of “growing” and “shrinking” objects. In half of the trials, the object also changed identity when the size change happened. According to the temporal contiguity hypothesis, and studies with a similar paradigm in monkeys, this training process should alter size tolerance. After the training phase, BOLD responses to each of the object images were measured in the scanner. Neural patterns in LOC and V1 contained information on size, similarity and identity. In LOC, the representation of object identity was partially invariant to changes in size. However, temporal contiguity training did not affect size tolerance in LOC. Size tolerance in human object-selective cortex is more robust to variations in input statistics than expected based on prior work in monkeys supporting the temporal contiguity hypothesis.

## 1 Introduction

Object recognition plays an important role in daily life. While humans are remarkably skilled at reliably identifying objects without much effort, this perceptual task poses a significant challenge to the visual system. Accurate object recognition is both selective and tolerant. On the one hand, object representations should be selective enough to ensure that objects with a highly similar appearance can be distinguished from one another. On the other hand, object representations should also be sufficiently tolerant to changes in appearance that are caused by the same object being seen under different viewing conditions (e.g. differences in viewing angle, distance, or lighting).

Learning has been shown to increase the selectivity of object recognition in human behavior (e.g., Folstein, Gauthier, & Palmeri, 2012; Goldstone, 1994, 1998), in the human brain (e.g., Brants et al., 2016; Folstein, Palmeri, & Gauthier, 2013; Freedman et al., 2006; Gillebert et al., 2009), and the monkey brain (e.g., Baker, Behrmann, & Olson, 2002; Kobatake, Wang, & Tanaka, 1998). Researchers believe that learning also boosts tolerance to changes in viewing conditions. Behavioral evidence suggests that invariant object recognition improves with age from infancy into adolescence and adulthood. The emergence of tolerance to different transformations has been shown to follow different developmental trajectories. A recent study on the neural basis of invariance has corroborated this finding, showing that size invariance develops earlier than view invariance (Nishimura et al., 2015).

There are multiple hypotheses about how tolerance might be learned. Here we investigate the so-called temporal contiguity hypothesis, which refers to the statistical properties of natural visual experience. In a natural context, images that quickly succeed each other and share visual features are more likely to represent different views of the same object rather than different objects. In other words, temporal contiguity is a useful cue that two images are of the same object. The visual system could take advantage of these natural statistics and learn to associate temporally contiguous images to link together multiple images of the same object into one transformation-tolerant object representation (Földiák, 1991; Rolls, 1992; Wallis & Rolls, 1997). The temporal contiguity hypothesis can be tested by manipulating the statistics of the visual input presented to participants. If the hypothesis is correct, it should not only be possible to increase naturally occurring tolerance by consistently presenting different images of the same object in rapid sequences, but also to break down existing tolerance and eventually build “incorrect” tolerance by presenting different objects together in the same way. In other words, presenting images in a temporally contiguous way should suffice to increase the representational similarity of these images, whether or not they appear together in the real world. Behavioral experiments in humans as well as single neuron studies in monkeys have employed this line of reasoning, and have provided evidence consistent with the temporal contiguity hypothesis.

Cox et al. (2005) provided behavioral evidence for temporal contiguity learning in humans for position invariance of objects. Participants were asked to fixate and then make a saccade toward an unfamiliar object (“greeble”) that appeared in the periphery. In half of the trials, unbeknownst to participants, the peripheral object was swapped for a different similar-looking object mid-saccade. Afterwards, in a same-different task, participants more often confused object pairs that had been swapped across retinal positions. These results suggest that different objects that were paired in a temporally contiguous way became more likely to be viewed as the same object (Cox et al., 2005). Similar behavioral effects have been reported for invariance to rotation in plane, rotation in depth, and illumination in face recognition (Wallis et al., 2009; Wallis & Bülthoff, 2001). In a recent study we used a design similar to the viewpoint tolerance experiment of Wallis et al. (2009) and also adapted it for an fMRI adaptation experiment. Both the behavioral and neuroimaging experiment provided evidence in favor of the temporal contiguity hypothesis (Van Meel & Op de Beeck, 2018). In all this human research, the confusions involved highly similar objects, such as different greebles or different faces, and effect size is relatively small.

In monkeys similar paradigms have resulted in strikingly strong effects on the responses of neurons in macaque inferior temporal cortex (Li & DiCarlo, 2008; 2010; 2012), resulting in mixing up the representations of highly dissimilar objects, such as for example a boat and a ball. In these studies, neurons’ responses to two objects that elicit strong (preferred object “P”) and moderate (non-preferred object “N”) responses were measured, before and after the monkeys were exposed to an altered visual world. During exposure phases, objects P and N were presented intermittently at different peripheral retinal positions (Li & DiCarlo, 2008) or in different sizes (Li & DiCarlo, 2010; 2012). The objects then changed retinal position (due to the monkeys making a saccade toward it; upwards or downwards) or size (bigger or smaller). Changes in one direction were accompanied by a change or ‘swap’ in object identity (e.g. P is replaced by N), while in the other case the object identity remained the same. In the test phases, responses to P and N were measured at all three positions or sizes, so that the tolerance of object selectivity (responses to P-N) could be assessed. The results of these experiments revealed that tolerance decreased significantly only at the swap position/size after just ∼1 hour of exposure (Li & DiCarlo, 2008; 2010), and that increased exposure up to ∼2 hours could even reverse object selectivity (Li & DiCarlo, 2010). Moreover, these effects seem to occur independent of the size and timing of reward (Li & DiCarlo, 2012).

Currently, there is a large difference between the human and monkey literature. All behavioral work in humans as well as our own recent fMRI study (Van Meel & Op de Beeck, 2018) have described modulations by temporal contiguity of the discriminability of very small differences between faces or objects, and have reported relatively small effect sizes. This stands in stark contrast with the electrophysiological animal work, where visual differences between objects are much larger, and where surprisingly large effects (cf. reversal of object preference) have been found. One potential reason for the discrepancy is that the human studies were typically inferring changes in representations indirectly from behavioral studies in which the sought-after change would actually impede performance. Possibly, large effects are confined to a situation in which a participant is not confronted with the consequences of the confusing representations that are induced by manipulations of temporal contiguity. For that reason we implemented a human study which was modeled as much as possible as the earlier monkey electrophysiology study focusing upon size invariance (Li & DiCarlo, 2010), using multi-voxel pattern analyses to measure neural representations. This study is the very first to use easily distinguishable objects as stimuli to investigate the temporal contiguity hypothesis in humans, and thus the first to use a design similar to the electrophysiological studies. Therefore, it provides an unprecedented test of the generalizability of the previously reported behavioral as well as neural effects.

## 2 Materials and Methods

### 2.1 Participants

Twenty-four volunteers (aged 20 to 37 years; mean age 23.9; seven male) participated in the experiment and were paid for their time. All participants had normal or corrected to normal vision, had no history of neurological or psychological problems, and were right-handed. Participants gave written consent at the start of the experiment. One participant (aged 24, female) was excluded because of steep drops in performance in multiple blocks of the training phase as well as the last four experimental runs in the scanner, suggesting sleepiness and lack of attention. We thus analyzed the data of 23 participants. This study was approved by the ethical committee of the Faculty of Psychology and Educational Sciences and the Medical Ethical Committee of the KU Leuven.

### 2.2 Stimuli

The stimuli were greyscale images of 12 different objects, arranged in six pairs (Figure 1).

**Figure 1.**
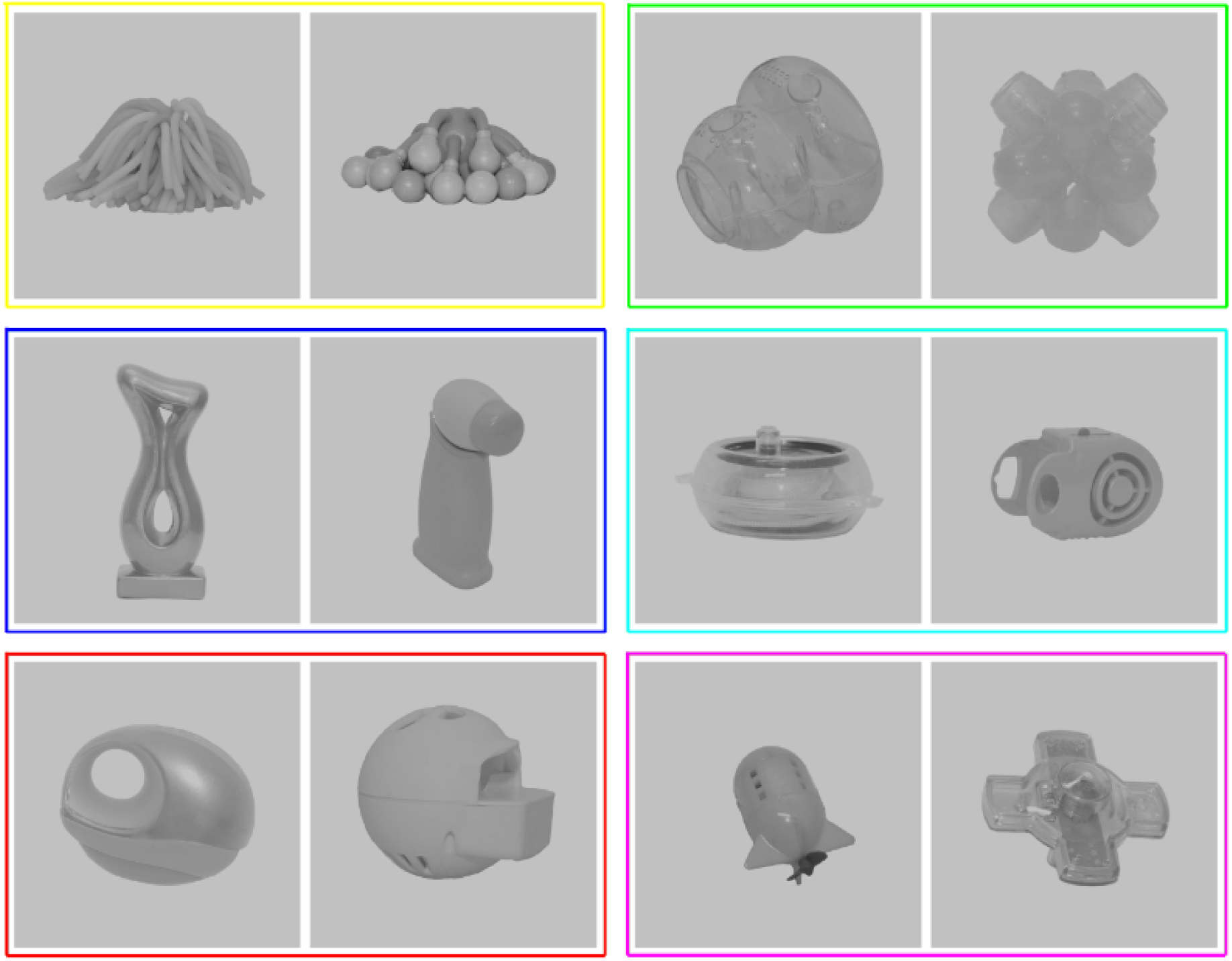
The stimulus set comprised 12 objects arranged in six pairs, with pairs consisting of similar but distinguishable objects. Pairs are framed in different colors for illustration purposes only.

The Novel Object and Unusual Name (NOUN) Database (Horst & Hout, 2016; downloaded from http://michaelhout.com/?page_id=759 on 20/02/2017) was used as a starting point for stimulus selection. All object images of the NOUN stimulus set were converted to greyscale and given a grey background. Images were additionally edited to homogenize luminance across the image set.

We then narrowed the stimulus set down to six pairs of objects, with pairs consisting of similar but distinguishable objects. To do this, we did not only take into account the subjective similarity of the objects, as reported in the NOUN Sorting Tables (also freely available at http://michaelhout.com/?page_id=759), but also computed the pixel-wise dissimilarity between all pairs of greyscale images. A first selection of pairs that were within the top 30 for pixel similarity and within the first quartile for subjective similarity resulted in a smaller set of 17 possible pairs. When multiple pairs included the same object, we only selected one of those pairs for the final stimulus set. We then excluded one pair that was an outlier in subjective similarity and two pairs for which the objects were not easily distinguishable as different objects to end up with our intended stimulus set size of six pairs.

### 2.3 Imaging parameters

Imaging data were collected on a 3T Philips Ingenia CX scanner with a 32-channel head coil at the Department of Radiology of KU Leuven. A multiband (MB 2) EPI imaging sequence with a TR of 2 s, TE of 30 ms, flip angle of 80°, and a 108 x 106 acquisition matrix was used for the functional images of the experimental runs. Each run consisted of 158 volumes (preceded by 2 dummy scans) containing 54 axial slices with a resolution of 2 x 2.04 x 2 mm and no interslice gap, providing whole-brain coverage. For the localizer runs, we used an EPI imaging sequence (no multiband) with a TR of 3 s, TE of 30 ms, flip angle of 90°, and a 104 × 106 matrix. Per localizer run, we obtained 75 volumes (preceded by 2 dummy scans) consisting of 47 axial slices of 2 mm thick with an in-slice resolution of 2.02 by 1.98 mm, and a gap of 0.2 mm. We also acquired anatomical images for all subjects, using an MP-RAGE sequence, with a voxel size of 1 × 1 × 1 mm.

### 2.4 Experimental procedure

The experiment consisted of a training phase of 1 hour outside the scanner, immediately followed by an fMRI scan session comprising eight experimental runs, three localizer runs and an anatomical scan. One extra experimental run was acquired in two subjects, and one extra localizer run was acquired in three subjects. For the training phase, stimulus presentation was controlled with a Dell laptop (Latitude 5480) running Windows 10 on a 14 inch monitor with a resolution of 1920 × 1080 and a frame rate of 60 Hz. Participants sat with their chins in a chin rest that was fixated to the table, so that the distance between their eyes and the center of the screen was held constant at 60 cm. In the scanner, pictures were projected onto a screen and were viewed through a mirror mounted on the head coil. MATLAB (Mathworks, inc.) and the Psychophysics Toolbox 3 (Brainard, 1997) were used to program the stimulus presentation in both phases.

#### 2.4.1 Behavioral training

On each trial, two images of different sizes were presented sequentially for 100 ms each, with the second image replacing the previous one, giving the impression of an object either growing or shrinking in size (Figure 2). Each trial was followed by an inter-trial interval of 800 ms. The first image of each trial was presented with a size of 7° of visual angle. The size of the second image was 11° in “growing” trials, and 3.5° in “shrinking” trials. A black fixation dot was presented in the center of the screen at all times.

**Figure 2.**
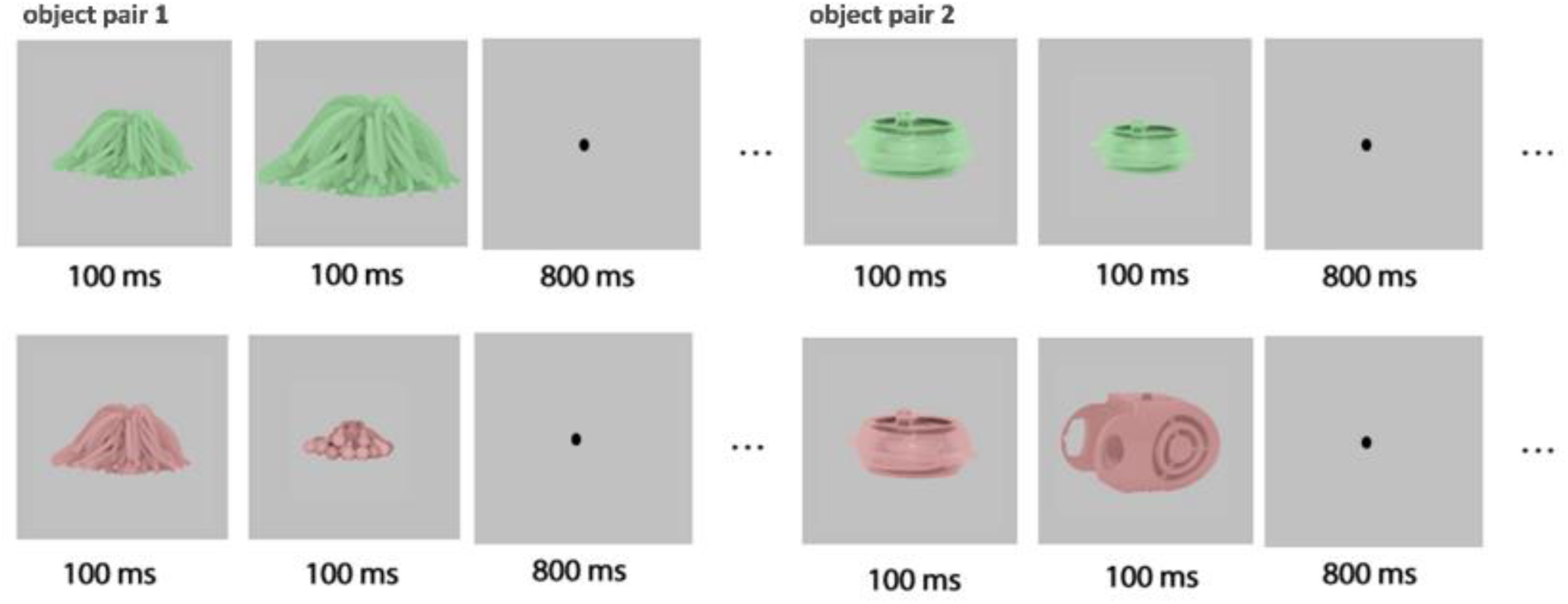
Example training trials for two out of four trained object pairs. For the pair on the left, the identity swap happens on shrinking, but not growing trials. For the pair on the right, the object identity swaps when growing, but not when shrinking. Across trials, each condition was shown in both green and red.

Two different types of trials were randomly interleaved during the training phase. In non-swap trials, the two images in a trial depicted the same object in two different sizes. In other words, these trials followed the natural image dynamics that we encounter in everyday environments. In the other half of the trials, the image dynamics were altered. In these so called swap trials, the images depicted two different object identities, with the first object being swapped for the other object of its pair when the change in size occurred. For two of the object pairs, further referred to as the “big-swap pairs”, the object was swapped for its partner object in growing trials, but the object identity remained unchanged in shrinking trials. For two other object pairs, further referred to as the “small-swap pairs”, the swap occurred on shrinking trials, but not on growing trials. The remaining two object pairs were not presented during the training phase, but served as untrained control condition in the scan session. The assignment of object pairs to the big-swap, small-swap and untrained conditions was counterbalanced across participants.

The training phase consisted of 10 training blocks of 320 trials each. Within each block, every object pair and every type of trial (shrinking/growing, swap/non-swap) was presented an equal number of times. The training lasted approximately an hour, with a total of 800 trials (400 swap and 400 non-swap trials) per object pair, and happened right before the start of the fMRI session. The number of trials per object pair is in line with Li and DiCarlo (2008). We presented one additional training block in the scanner as a short training refresher. This training refresher was presented during the anatomical scan, which took place after half of the experimental runs were collected.

During training, participants were faced with an orthogonal task. The stimuli randomly appeared in red or green, and participants were required to indicate the color of the current trial by pressing one of two buttons. An equal number of red and green trials was presented, and task difficulty increased by gradually fading the colors to grey in four interleaved one-up two-down adaptive staircases.

#### 2.4.2 Experimental fMRI runs

During the experimental runs, all twelve objects (i.e. eight trained objects and four control objects) were presented one by one in all three sizes in an event-related design. On each trial, an object image was shown for 1.5 s, followed by a fixation point for 1.5 s. Over the course of a run, each object was presented twice (once in the first half, and once in the second half of the run) in each of the three sizes. We interspersed extra fixation trials (25%) of 1.5 s at random time points. A fixation point was presented for 14 s prior to the first stimulus presentation and after the last stimulus. The orthogonal task was the same as during training.

#### 2.4.3 Localizer fMRI runs

During the localizer runs, grayscale pictures of faces, non-living objects and scrambled textures were presented in a blocked design. Blocks contained 20 images of one category (faces/objects/scrambled) which were presented for 300 ms each, with blanks of 450 ms between images. Blocks lasted 15 s. A fixation block was presented at the beginning, middle and end of each run. Each category was presented four times, in pseudo-random order (twice before and twice after the middle fixation block). Participants performed a one-back repetition detection task by pressing a button when the same picture was presented two times in a row. Three repetitions were presented per block.

## 3 Data Analyses

### 3.1 Preprocessing and General Linear Model

The Statistical Parametric Mapping software package (SPM12, Wellcome Department of Cognitive Neurology, London) was used to preprocess the imaging data of all functional runs.

Functional images of all runs were slice time corrected, spatially realigned to correct for head motion, and coregistered to the anatomical images. All images were then normalized to MNI space. During normalization, functional images were re-sampled to a voxel size of 2 × 2 × 2 mm. Finally, functional images were spatially smoothed (4 mm FWHM). Head motion was defined as excessive if it exceeded 2 mm between two adjacent time points within a run. Excessive head motion resulted in the exclusion of one experimental run in two subjects, and three experimental runs in one subject. In two subjects, one localizer run was excluded because of an issue with the timing of stimulus presentation.

The preprocessed data was modeled for each voxel, for each run, and for each participant using a General Linear Model (GLM). The GLM of the experimental runs included 36 regressors for the experimental conditions (i.e. twelve objects in three sizes), and six regressors for the motion correction parameters (*x,y*,z for translation and rotation, extracted during preprocessing). For the localizer runs, the GLM consisted of three regressors for the stimulus conditions (faces, objects, and scrambled images) and six motion correction regressors.

### 3.2 Regions of interest

Multiple regions were chosen as region of interest (ROI; Figure 3). Given its important role in object recognition, the lateral occipital complex (LOC) was the main region of interest. The fusiform face area (FFA) and V1 were chosen as a high-level and low-level visual control region, respectively, which could speak about the specificity of potential learning effects in LOC. Finally, we defined two ROIs involved in associative learning.

**Figure 3.**
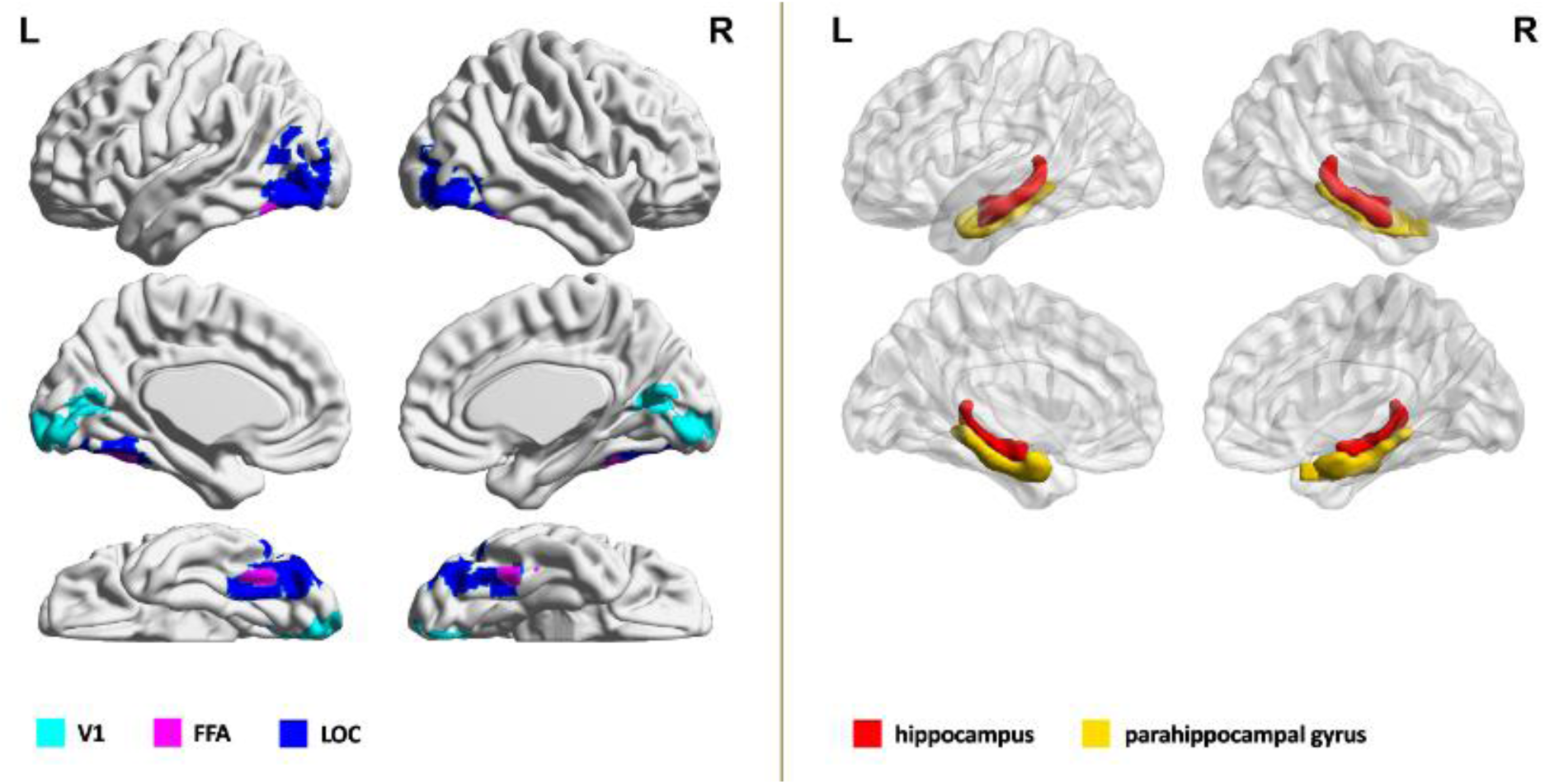
Regions of interest (ROI) for one subject. Visual ROIs are presented in the left panel, and medial temporal lobe (MTL) ROIs are presented on the right. This figure was made using BrainNet Viewer (Xia, Wang, & He, 2013). The visual ROIs were projected on the surface of the ICBM152 template. The MTL ROIs were drawn as volumes in the same template.

The high-level object- and face-selective visual regions LOC and FFA were defined using functional data of the localizer runs. Subject-specific functional activation maps for the objects scrambled and faces > objects contrasts were thresholded at p = 0.0001 and masked with the LOC and FFA parcels of Julian et al. (2012), respectively. The threshold for a contrast was lowered in case both hemispheres contained less than 20 active voxels. This resulted in a more lenient threshold for LOC in one subject (p = 0.01). For one subject, we could not define FFA even at p = 0.01. Low-level region V1 was defined using the Automated Anatomical Labeling (AAL) atlas (Tzourio-Mazoyer et al., 2002) combined with functional data of the experimental runs. More specifically, subject-specific functional activity maps for the contrast of all stimuli >fixation were thresholded at p = 0.0001 and masked with the AAL’s calcarine region (not dilated). Stimulation in the localizer runs was less representative for the visual experience in the experiment in terms of stimulus size and size variation. Therefore, we used data of the experimental rather than the localizer runs to define retinotopic area V1.

The medial temporal lobe (MTL) has been shown to be involved in associative learning (e.g. (Schapiro, Kustner, & Turk-Browne, 2012; Schendan et al., 2003; Turk-Browne et al., 2010). While the motivation for our study was derived from the visual learning literature, the paradigm could also be rephrased as an associative learning paradigm in which swap trials would cause the buildup of an association between two previously unfamiliar objects. Hence, we selected the hippocampus and parahippocampal gyrus as ROIs in addition to the visual ROIs. To define these regions, we used the AAL masks (hippocampus and parahippocampal; not dilated) without functional constraints. Note that the imaging parameters and design were not optimized for MTL coverage, hampering a more precise delineation of the hippocampus and parahippocampal gyrus and their subregions.

### 3.3 Neural dissimilarity matrices

For each ROI in each subject, a neural dissimilarity matrix was constructed from the beta weights (estimated in the GLM) of that ROI’s voxels for each of the 36 stimulus conditions. We used the cross-validated Mahalanobis distance (Walther et al., 2016) as a measure of neural dissimilarity. This measure is the cross-validated Euclidean distance normalized by the covariance of the training sample, and is also called the linear discriminant contrast (LDC) as it is directly related to linear discriminant analysis (LDA). The training sample covariance matrices were regularized using the oracle approximating shrinkage estimator (Chen, Wiesel, Eldar, & Hero, 2010) to deal with potential rank deficiency (Walther et al., 2016). We used a leave-one-run-out cross-validation procedure. The 36 × 36 neural dissimilarity matrices represented all pairwise distances between conditions, averaged over cross-validation folds. We used code written by J. Brendan Ritchie (Ritchie & Op de Beeck, 2019) for computation of the LDC, combined with the CoSMoMVPA toolbox (Oosterhof, Connolly, & Haxby, 2016) to set up the cross-validation procedure.

### 3.4 The effect of temporal contiguity training on size tolerance

For each subject and ROI, distances of interest were calculated by averaging across the relevant cells of the dissimilarity matrix. More specifically, LDC distances between images of objects within a pair differing in size and/or identity (Figure 4) were extracted and averaged across pairs of the same type (i.e. big-swap pairs, small-swap pairs, control pairs). We then averaged each distance across all experimental pairs (i.e. big-swap and small-swap) to even out any effect of object size. For experimental pairs, distances A_swap_ and B_swap_ always refer to distances between the medium and swap size, and distances A_non-swap_ and B_non-swap_ always refer to distances between the medium and non-swap size. Given that control pairs were not presented during training, we cannot denote a “swap size” and “non-swap size” for those pairs. For control pairs, distances receive the subscript “control” and are averaged across sizes (e.g. A_control_ is the mean of distances A_medium-to-big_ and A_medium-to-small_). We tested whether distances were significantly larger than zero using one-tailed one sample t-tests.

**Figure 4.**
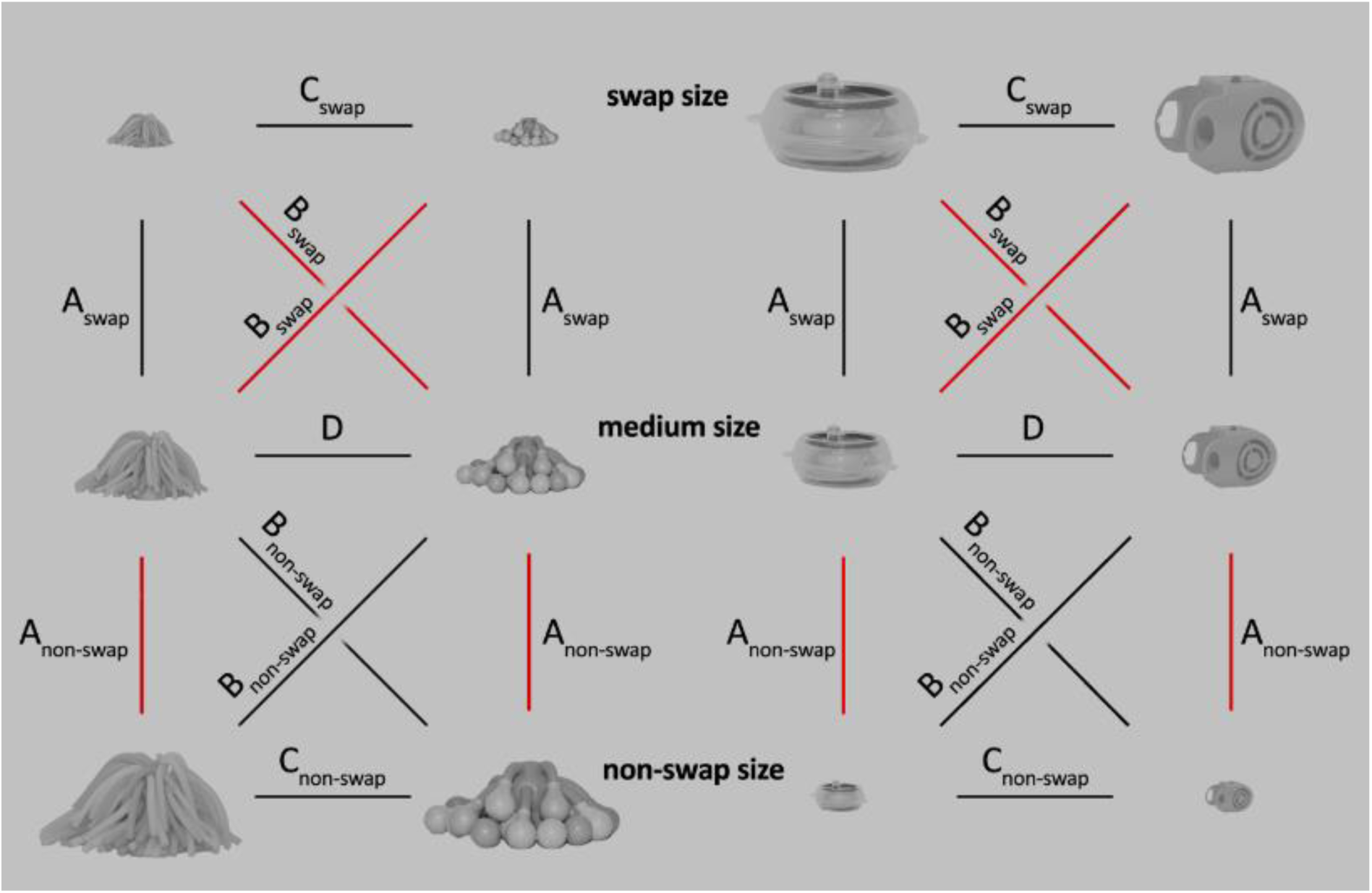
Within-pair distances for two experimental pairs. In this example, a small-swap pair is shown on the left, and a big-swap pair is shown on the right. Distances A and B were of interest to test the effect of temporal contiguity training. The distances connecting stimuli that were associated during training are shown in red. For control pairs (not pictured), the subscript “control” was used instead of “swap” and “non-swap”, and distances were averaged across sizes.

We first tested whether the ROIs represent objects in a size tolerant way, using data from the control pairs as these are not influenced by any training effects. If object representations are tolerant to changes in size, neural distances to images of the same object in a different size should be smaller than neural distances to images of a different object in a different size. We thus tested whether there was a difference between distance A_control_ (distance to same object) and distance B_control_ (distance to other object of the pair) using two-tailed paired t-tests.

An effect of the training can reveal itself in different ways. On the one hand, the medium size image (7° of visual angle) of each object was consistently followed by an image of that same object in the non-swap size (Figure 2, upper row). This process mimics natural image dynamics and could, according to the temporal contiguity hypothesis, result in increased size tolerance. In other words, these image dynamics could strengthen the link and thus decrease the neural distance between the medium and non-swap size images of that object (distance A_non-swap_). At the same time, this process could reinforce object selectivity and thus increase the distance to the non-swap size image of the other object of its pair (distance B_non-swap_). On the other hand, the medium size image of each object was also consistently followed by an image of the other object of its pair in the swap size (Figure 2, lower row). If the temporal contiguity hypothesis is correct, the training could thus also strengthen the association between the medium size image of an object and the swap size image of its counterpart and build new incorrect size tolerance (decreased distance B_swap_). At the same time, it could “break” the correct size tolerance for that size and increase the neural distance between the medium and swap size images of that object (distance A_swap_).

We tested whether there is a difference between distances A_swap_ and A_non-swap_ (distance to same object) and between distances B_swap_ and B_non-swap_ (distance to other object) in the experimental pairs, using two-tailed paired t-tests. A positive difference between distances A_swap_ and A_non-swap_ can be due to an increase in distance A_swap_ a decrease in distance A_non-swap_, or both. A negative difference between distances B_swap_ and B_non-swap_ can be due to a decrease in distance B_swap_, an increase in distance B_non-swap_, or both. Together, these tests can thus account for all possibilities mentioned above.

Apart from testing the temporal contiguity hypothesis in the experimental pairs, we also tested whether the distances of interest differed between experimental pairs and control pairs, using two-tailed paired t-tests. Distances A_swap_ and A_non-swap_ were each tested against distance A_control_, and distances B_swap_ and B_non-swap_ were tested against distance B_control_. Positive differences for A_swap_ and B_non-swap_ and negative differences for A_non-swap_ and B_swap_ would be in line with the temporal contiguity hypothesis.

We used an α-level of 0.05 for all tests and applied a Bonferroni correction for the number of ROIs (five), resulting in a corrected α-level of 0.01. Note that this approach is rather conservative, because our ROIs differ in status: we chose one main ROI (LOC), two visual control regions (V1 and FFA), and two regions which were selected post-hoc given their relevance to associative learning (hippocampus and parahippocampal gyrus). All reported p-values are uncorrected, and thus considered as significant if p < 0.01.

### 3.5 The representation of object similarity and size

Due to the way in which we selected stimulus pairs (see 2.2 Stimuli), objects within a pair are more similar to each other, both in pixel-wise similarity and subjective similarity, than objects belonging to different pairs. During the experimental fMRI runs, all object images were presented in three different sizes. Object images of different sizes can thus either differ one size step (small vs. medium, or medium vs. big) or two size steps (small vs. big) from each other. We tested whether these properties of our stimulus set were represented in the neural activation patterns in each of the ROIs. For the purpose of these analyses, data was collapsed across all pairs irrespective of training status. To test the effect of object similarity, we calculated a within-pair and between-pair measure for each participant. For the within-pair measure, we calculated the mean of all possible pairwise distances between images within a pair. This includes all distances A to D, plus the distances between small and big images within a pair. We then averaged that measure across all pairs. The between-pair measure is the mean of all possible pairwise distances between all images of the stimulus set, excluding the within-pair distances. The within-pair and between-pair measure were then contrasted using a two-tailed paired t-test (corrected α = 0.01). For each participant, we also calculated a one size step measure and a two size step measure. These measures were calculated as the mean of all possible pairwise distances between images differing one or two size steps respectively, irrespective of the object identity. We contrasted the one size step and two size step measures using a two-tailed paired t-test (corrected α = 0.01).

For ROIs that showed an effect of the degree of size difference and object similarity, we additionally calculated two measures per participant intended to map the relative contribution of object identity and size to the ROIs neural representations, and compare it between ROIs. For the object discrimination measure, we took the pairwise distances between different objects of a pair in the same size (distances C and D) and averaged those across sizes and across pairs. Note that this measure was calculated within pairs, instead of between each object and all other objects. Given the high object similarity within pairs, it thus provides a strict test of the differentiation of object identity. The size discrimination measure was the average of all pairwise distances between images of the same object differing one size step from each other (distances A). Distances spanning two size steps were left out to ensure that our size discrimination measure was as strict as possible. A two-tailed paired t-test was used to compare size and object discrimination measures within ROIs (tested in LOC and V1; corrected α = 0.025).

To compare the relative contribution of object identity and size between ROIs, we first normalized the difference between size and object discrimination by the absolute value of their sum. We then compared this new ratio between ROIs using a two-tailed paired t-test (α = 0.05).

## 4 Results

### 4.1 Within-pair images can be distinguished reliably by LOC and V1

We first checked within-pair discrimination, because it is a necessary condition to further investigate how training might change the representation of the images in a pair. In LOC, within-pair LDC distances A to D were all significantly larger than zero in the experimental pairs (A_swap_: t(22) = 2.92, p = 0.004; A_non-swap_: t(22) = 5.16, p = 1.79e-5; B_swap_: t(22) = 5.27, p = 1.37e-5; B_non-swap_: t(22) = 5.75, p = 4.38e-6; C_swap_: t(22) = 3.09, p = 0.0027; C_non-swap_: t(22) = 3.94, p = 0.0003; D: t(22) = 3.09, p = 0.0027; ; one-tailed one sample t-tests). In the control pairs, all distances were significantly larger than zero as well (A_control_: t(22) = 4.11, p = 0.0002; B_control_: t(22) = 5.14, p = 1.86e-5; C_control_: t(22) = 3.10, p = 0.0026; D: t(22) = 2.76, p = 0.0057; one-tailed one sample t-tests). These results indicate that, overall, neural patterns for different images within a pair could reliably be distinguished in LOC (Figure 5).

**Figure 5.**
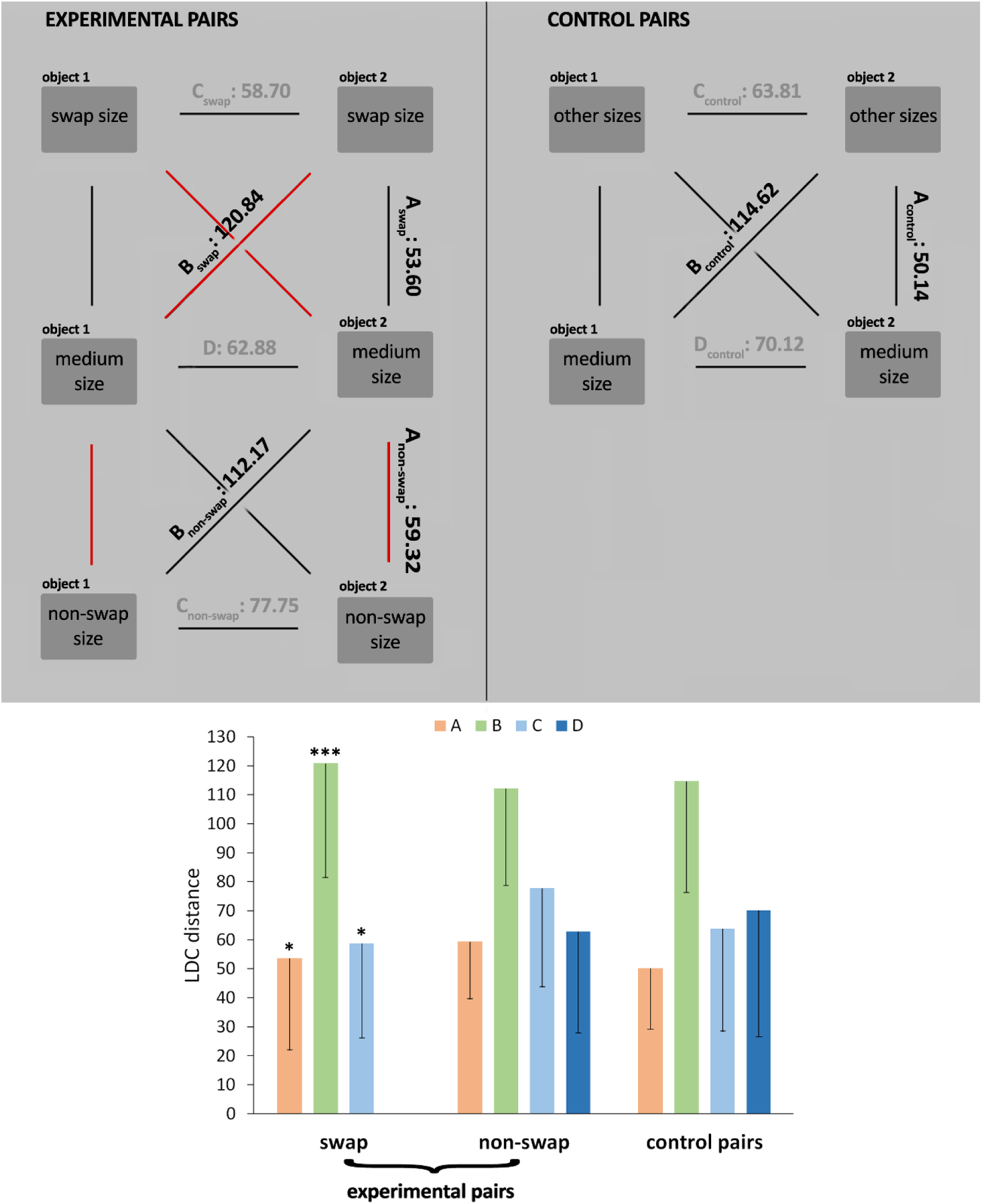
Within-pair distances in LOC across objects and participants. The upper panel shows a schematic overview of the mean distances in experimental pairs and in control pairs. Distances used to analyse the training effect are printed in black. Distances not used in those analyses are printed in grey. As in Figure 4, red lines connect stimuli that were associated during training The lower panel shows a bar plot of the same mean distances. Error bars represent one-sided 95% confidence intervals. * p < 0.01, ** p < 0.001, *** p < 0.0001

Neural patterns for the different images within a pair were also reliably distinct in V1 (all t(22) >3.08, p < 0.0027; one-tailed one sample t-tests). None of the other control regions could make the distinction between images in a pair (FFA: all t(21) < 2.19, all p > 0.0199; hippocampus: all t(22) < 1.97, all p > 0.0305; parahippocampal gyrus: all t(22) < 1.48, all p > 0.0774; one-tailed one sample t-tests).

### 4.2 Temporal contiguity training does not affect size tolerance

In LOC, distances to images of the same object in a different size (A_control_) differed significantly from distances to images of a different object in a different size (B_control_), with the former being smaller (t(22) = -4.62, p = 0.0001; two-tailed paired t-test). Neural representations of object identity in LOC are thus tolerant to changes in size. This was not the case in the other ROIs (all |t| < 1.59, all p > 0.1272; two-tailed paired t-tests).

Contrary to the predictions from the temporal contiguity hypothesis, the training did not induce any difference in size tolerance between the swap size and the non-swap size (A_swap_ vs. A_non-swap_: t(22) = -0.44, p = 0.6673; B_swap_ vs. B_non-swap_: t(22) = 0.73, p = 0.4744; two-tailed paired t-tests). Moreover, the distances of interest in LOC did not differ between experimental pairs and control pairs (A_swap_ vs. A_control_: t(22) = 0.20, p = 0.8466; A_non-swap_ vs. A_contro_l: t(22) = 0.85, p = 0.4018; B_swap_ vs. B_control_: t(22) = 0.40, p = 0.6942; B_non-swap_ vs. B_control_: t(22) = -0.19, p = 0.8498; two-tailed paired t-tests). In other words, we did not find any evidence that temporal contiguity training can build or break size tolerance in LOC (Figure 6).

**Figure 6.**
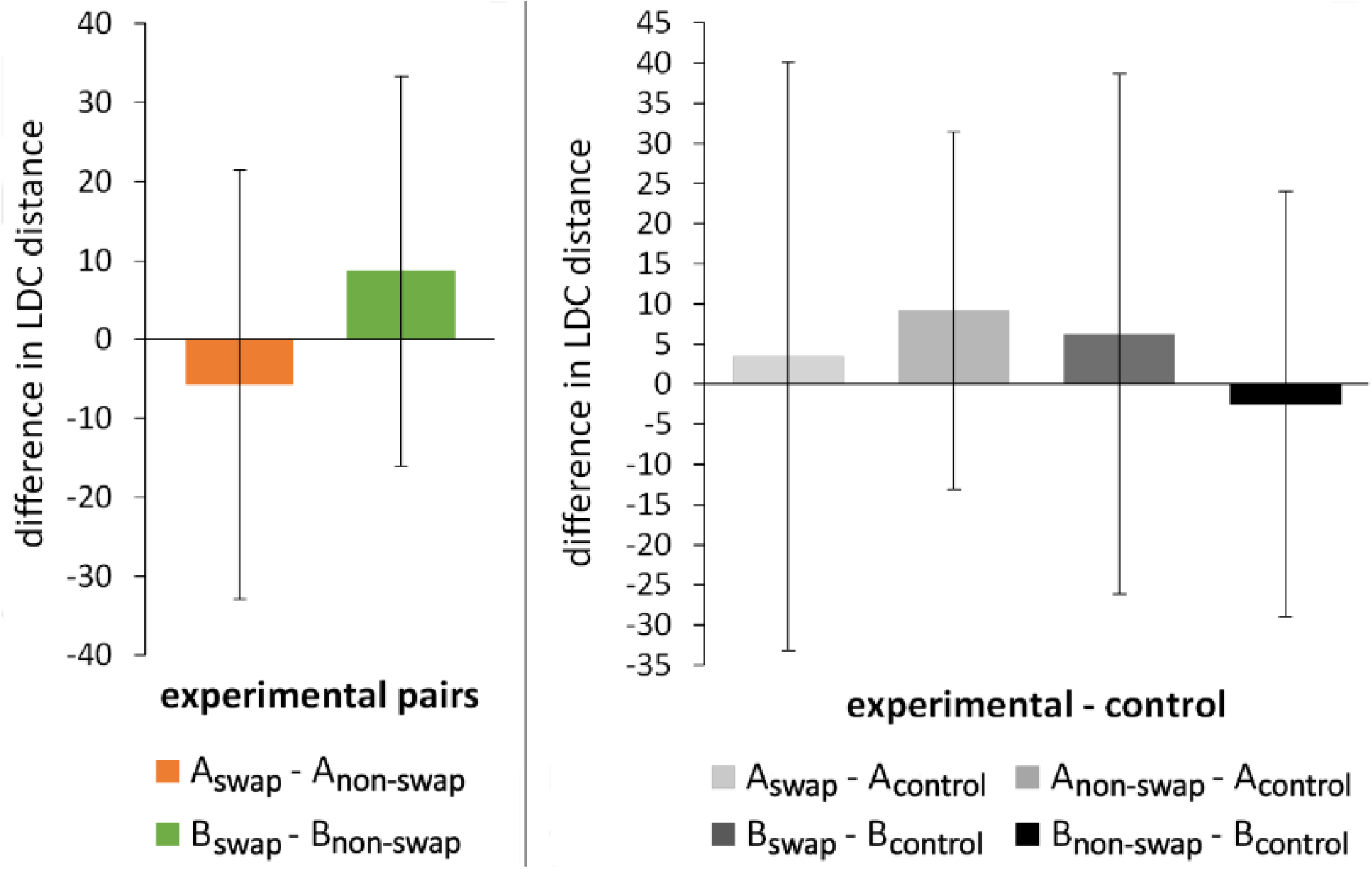
The effect of training on distances of interest in LOC. The left panel shows the difference between distance A_swap_ and A_non-swap_ and between distance B_swap_ and B_non-swap_ in the experimental pairs. The right panel shows differences between experimental pairs and control pairs. More specifically, distances of interest in the experimental pairs are compared with their size-controlled equivalents in the control pairs (see text). Error bars denote 95% confidence intervals of the differences. All differences were non-significant.

The prior sections already indicate that we lack sensitivity in the other visual ROIs to pick up potential effects, nor did we predict any effects there. Thus it is not a surprise that the other visual ROIs also did not show any difference between distance A_swap_ and A_non-swap_ or between distance B_swap_ and B_non-swap_ in the experimental pairs (V1: both |t(22)| < 0.93, both p > 0.3651; FFA: both |t(21)| < 0.44, both p > 0.6611; two-tailed paired t-tests). Differences between the distances of interest in the experimental and control pairs did not reach significance either (V1: all |t(22)| < 0.93, all p > 0.3643; FFA: all |t(21)| < 1.83, all p > 0.0826; two-tailed paired t-tests).

We addressed two further considerations that are relevant given the null result of training in LOC. First, it is important to rule out that our experiment would just lack the sensitivity to find existing effects of training. Here it is relevant to emphasize that in LOC we were able to pick up size tolerance (by comparing distances A_control_ with B_control_) with very high significance (p = 0.0001). The size tolerance measured here has an effect size d_z_ of 0.9638 (i.e. t/vn = 4.6223/v23), and our experiment has a power of 0.9929 to detect an effect of this size with a two-tailed paired t-test and an α of 0.05 (calculated using G*Power 3.1; Faul, Erdfelder, Lang, & Buchner, 2009). In a study by Li and DiCarlo (2008) object selectivity at the swap position, or in other words position tolerance, was reduced by half after a similar number of exposure trials as used here (see their Figure 2a). Therefore, we calculated the power of our experiment to measure a reduction of the observed size tolerance by half (d_z_ = 0.4819). Using a two-tailed paired t-test and an α of 0.05, this power is 0.5986. The power further increases to 0.7238 when using a one-tailed test which would be appropriate given that the prediction is directional (reduction in size tolerance). Thus, it is not very likely that we miss an existing effect due to lack of sensitivity, unless the effect is smaller than expected based on prior research.

Second, the interpretation of the absence of any effect of temporal contiguity in the visual system might be constrained by potential effects in other brain regions that code for different aspects of the learning event. For this reason, we also included control ROIs known to be involved in associative learning. It is hard to predict exactly how our training protocol would influence representations in regions such as hippocampus and parahippocampal gyrus. Standard associative learning protocols include training with random pairings of objects, and find increased neural similarity for objects that have been paired. The critical difference in our paradigm is that object pairings are size-dependent, so that an object is paired with a different-size version of itself at the non-swap size (distance A_non-swap_), and with a different-size version of another object at the swap size (distance B_swap_). The reliability of neural patterns in these medial temporal lobe ROIs is low (see Section 4.1), and as such a null result in these ROIs would not be very informative, but nevertheless we checked whether there is any indication of a coding for these associations. This seemed to be the case for distance A_non-swap_. In both the hippocampus and parahippocampal gyrus, distance A_non-swap_ in the experimental pairs was smaller than distance A_control_ (hippocampus: t(22) = -2.20, p = 0.0386; parahippocampal: t(22) = -2.17, p = 0.0408; two-tailed paired t-tests). This finding suggests that training led to increased representational similarity (i.e. smaller neural distance) between the medium and non-swap size images of objects, and thus the temporal association between an object and its temporally associated different-size version was coded for in the medial temporal lobe. Note however that these effects do not survive our correction for multiple comparisons. No similar effect was noted for the temporally associated different object at the swap size (distance B_swap_), nor was there any effect for the other distances (hippocampus: both |t(22)| < 1.17 and p > 0.2560; parahippocampal: both |t(22)| < 1.18 and p > 0.2530). Comparing distances for swap and non-swap sizes within experimental pairs did not reveal any effects either (hippocampus: both t(22) < 1.76 and p > 0.0917; parahippocampal: both |t(22)| < 1.04 and p > 0.3080). Thus, there is some, albeit limited, evidence that the image associations built up during training are represented outside visual cortex.

### 4.3 Representation of similarity, size and object identity

Finally, we present a more detailed characterization of how representations change from V1 to LOC in terms of shape similarity and size invariance. Both LOC and V1 show a reliable representation of object similarity and size. The difference in similarity between all images within a pair and all images belonging to different pairs was picked up by LOC (t(22) = -6.63, p = 1.14e-6) and V1 (t(22) = -5.14, p = 3.75e-5), but not by the other ROIs (all |t| < 1.24, all p 0.2316; two-tailed paired t-tests; Figure 7, right panel, purple bars). The difference between distances covering one size step and two size steps (independent of object identity) was also represented in these regions (LOC: t(22) = -6.30, p = 2.43e-6; V1: t(22) = -5.66, p = 1.09e-5; other ROIs: all |t| < -1.94, all p > 0.0657; two-tailed paired t-tests; Figure 7, right panel, yellow bars).

**Figure 7.**
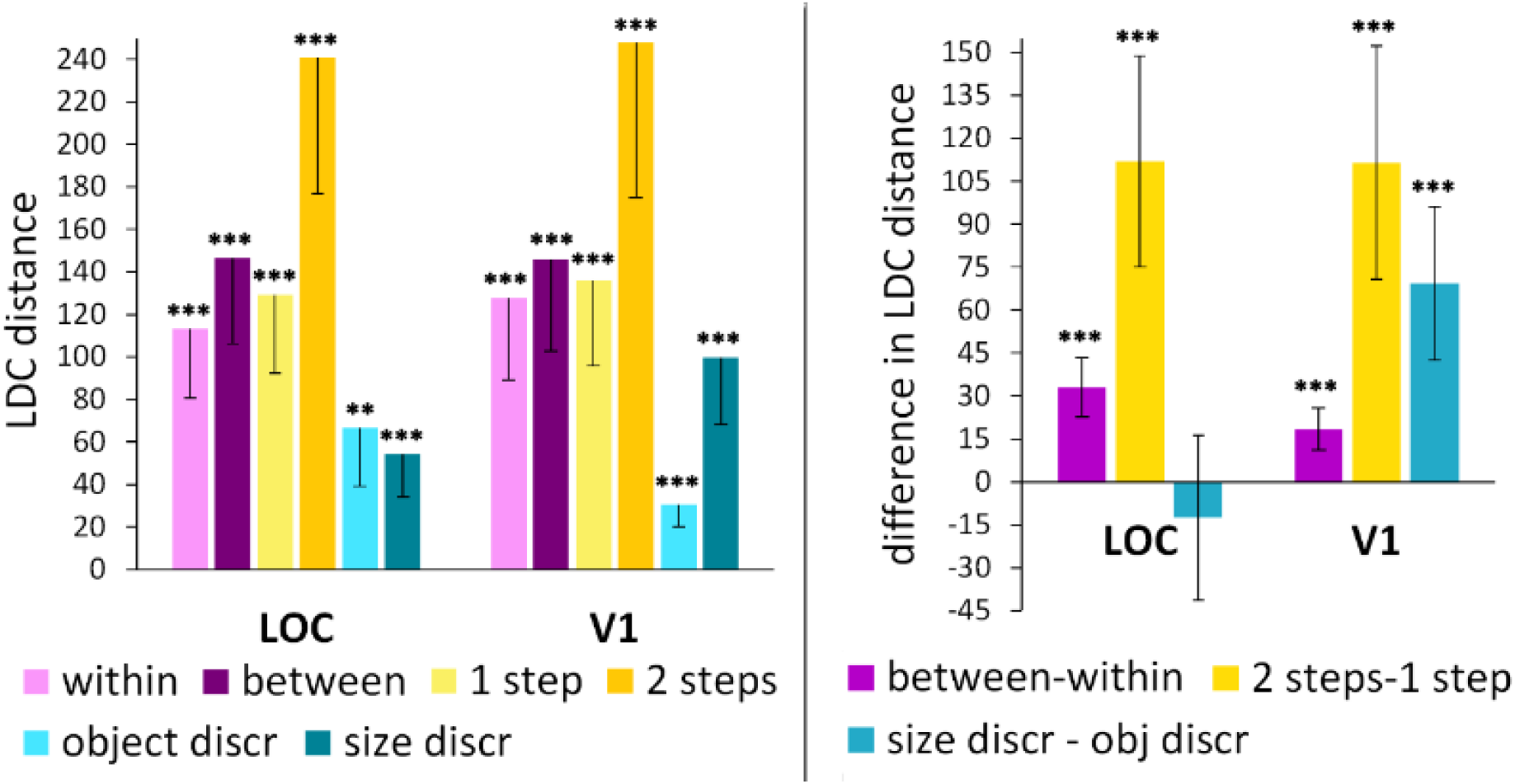
Effects of object similarity, size, and object identity in LOC and V1. The left panel shows the four measures that were calculated to test the effect of object similarity and size difference, as well as the object discrimination and size discrimination index. Error bars denote one-sided 95% confidence intervals. The right panel shows the effects of similarity and size, and the difference between size and object discrimination measures. Error bars represent 95% confidence intervals of the differences. ** p < 0.001, *** p < 0.0001

Neural representations in LOC and V1 seem to be rich in information: these regions are sensitive to differences in similarity and size, and can differentiate between individual images (see 4.1). While these results indicate similarities between the neural representations in LOC and V1, we might also expect certain differences, given the regions’ distinct location and function in the visual system. The object discrimination and size discrimination measures allowed us to investigate each region’s representations in more detail and to compare the relative contribution of object identity and size between regions.

In LOC, the neural distance between objects of the same pair and the same size (i.e. the object discrimination measure) was significantly larger than zero (t(22) = 4.13, p = 0.0002; one-tailed one sample t-test). The neural distance between images of the same object differing one size step (i.e. the size discrimination measure) was also significant (t(22) = 4.60, p = 6.94e-5; one-tailed one sample t-test; Figure 7, left panel, turquoise bars). There was no difference between the two measures (t(22) = -0.90, p = 0.38; two-tailed paired t-test; Figure 7, right panel, turquoise bar).

In V1, the object discrimination and size discrimination measures were both significant as well (object discrimination: t(22) = 5.03, p = 2.47e-5; size discrimination: t(22) = 5.45, p = 8.86e-6; one-tailed one sample t-tests; Figure 7, left panel, turquoise bars). Moreover, the size discrimination index was significantly larger than the object discrimination index (t(22) = 5.37, p = 2.14e-5; two-tailed paired t-test; Figure 7, right panel, turquoise bar).

While the absence of a difference in LOC and the existence of a difference in V1 already provides a suggestion that the relative contribution of size and object identity to the neural representations differs between the regions, we also tested this directly by comparing the normalized difference of the measures (i.e. ratio of the difference and the absolute value of the sum) between the regions. This test revealed a significant difference between V1 and LOC (t(22) = 2.96, p = 0.0073; two-tailed paired t-test; Figure 8, right panel).

**Figure 8.**
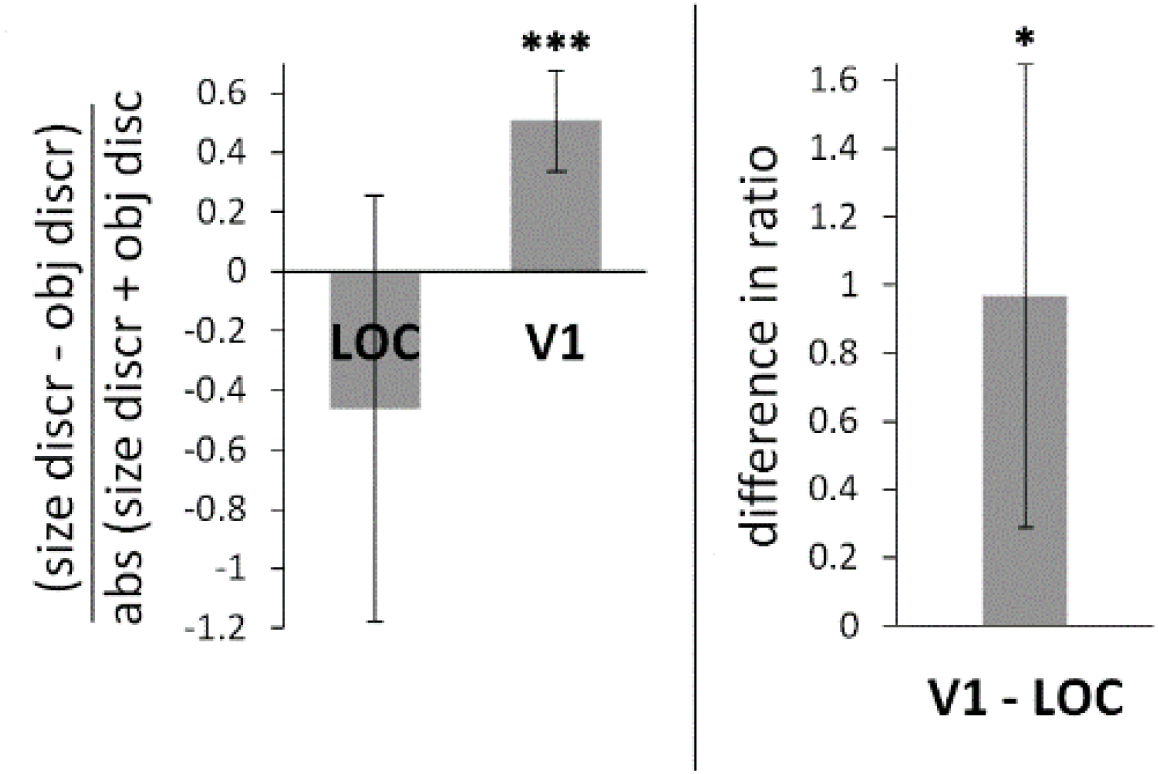
Relative contribution of identity and size representation in LOC and V1. The left panel shows the difference between size and object discrimination measures divided by the absolute value of their sum. Note that this ratio is the normalized versions of the turquoise bars in the right panel of Figure 7. The right panel shows the difference in this ratio between V1 and LOC. Error bars represent 95% confidence intervals. * p < 0.01, *** p < 0.0001

## 5 Discussion

In this study, we investigated neural object representations in multiple regions of the visual cortex and MTL using multivariate pattern analyses. In LOC and V1, different images of objects with a similar shape can be distinguished from one another. The representational similarity in these regions reflects a combination of information on size differences, degree of similarity, and object identity. Moreover, the representation of object identity in LOC is partially size invariant, and more size invariant than in V1, as such confirming what we know about information processing in these regions. The aim of this study was to investigate the effect of training protocols designed to influence this size invariance. We exposed participants to sequences of images of objects changing in size, and in half of the cases also changing identity. According to the temporal contiguity hypothesis, this process should alter size tolerance. However, we did not find any evidence that temporal contiguity training can build or break size tolerance in LOC.

The stimulus set used here was adapted from the Novel Object and Unusual Name (NOUN) Database. The NOUN Database consists of images of real 3D novel objects. Novelty of the objects was confirmed by self-report and a lack of consensus in naming and/or identifying the objects (Horst & Hout, 2016). Given that these objects are not instinctively associated with a certain name, changes in object identity happening during the training phase were less likely to be verbalized (e.g. “the boat changed into a fish”). We reasoned that verbalization of these identity changes could counteract effects of temporal contiguity learning in the swap conditions. We thus avoided that potential problem by using novel rather than familiar objects. Likewise, the subjective similarity and shape similarity of objects in a pair further helped to avoid verbalization. In the monkey studies by Li and DiCarlo (2008, 2010, 2012) stimuli were objects that are familiar to humans (e.g. a boat and a fish) but can be assumed to be novel to the monkeys. Using unfamiliar objects in this study thus had the additional advantage that prior experience with the stimuli is more comparable to the monkey studies.

Our finding of partial size tolerance of object discrimination in LOC is in agreement with electrophysiological evidence in monkeys indicating that only a subset of neurons in inferotemporal (IT) cortex show size-invariant responses, and that object identity as well as size information is represented in IT (Hung et al., 2005; Ito et al., 1995). Our results are also consistent with the findings of human neuroimaging studies by Eger et al. (2008) and Eger, Kell, & Kleinschmidt (2008). We showed that representational information in LOC is detailed enough to allow for discrimination of objects with similar shapes. While we did not use stimuli from well-known everyday categories, this finding translates well to the within-category object discrimination reported in Eger et al. (2008) and Eger, Kell, & Kleinschmidt (2008), given that objects within a category typically share more similar shape features than objects belonging to different categories. Our results also revealed that representational distances across two size steps were significantly larger than distances across one size step. This matches the graded size sensitivity found by Eger, Kell, & Kleinschmidt (2008).

We showed that, apart from LOC, the different object images could also be reliably discriminated by V1. The neural patterns in both LOC and V1 carried information about object identity as well as size, but their relative contribution differs between the regions, with LOC favoring identity information, and V1 favoring size information. Similar dissociations between V1 and LOC were reported by Eger et al. (2008) and Eger, Kell, & Kleinschmidt (2008). We found that in V1, object discrimination was not tolerant to changes in size, indicating that the object identity effect in V1 can be explained by pixelwise differences between objects. This raises the question to what extent the neural representations in LOC also reflect pixelwise similarity rather than a more abstract representation of object identity. However, the size tolerance in LOC and the aforementioned dissociation with V1 strongly suggest that the results in LOC are not purely driven by low-level similarity.

Contrary to the predictions of the temporal contiguity hypothesis, we did not find any changes in size tolerance in LOC as a consequence of the training protocol. Presenting object images together in a temporally contiguous way over the course of a 1 hour period did not alter the neural distance between the representations of these images. We did not find an effect when comparing neural distances between the swap and non-swap conditions within pairs, and neither when comparing neural distances between experimental pairs and control pairs. This lack of an effect of temporal contiguity training contrasts with results of previous studies in humans (Cox et al., 2005; Van Meel & Op de Beeck, 2018; Wallis et al., 2009; Wallis & Bülthoff, 2001). Apart from the fMRI experiment in our earlier study (Van Meel & Op de Beeck, 2018), all previous human studies have indirectly inferred changes in neural representations from changes in behavioral performance on a same/different task. When participants consciously try to discriminate between objects, extra attention might be drawn to the differences between those objects. Moreover, the binary same/difference decision required by the task might make it harder to capture subtle changes in representational similarity. As a result, behavioral measures might cause reduced effect sizes and cover up more substantial underlying changes brought about by temporal contiguity. Following this line of reasoning, studies using neural measures instead of behavioral ones should result in larger effects. Instead, we reported a null result. It thus seems unlikely that the discrepancy in results is caused simply by using a neural measure rather than a behavioral readout.

The absence of a temporal contiguity effect is also not in line with evidence from electrophysiological studies in primates (Li & DiCarlo, 2008; 2010; 2012). We studied human participants instead of macaque monkeys. Given the well documented anatomical and functional similarities between monkey IT and human LOC, this difference in studied species in itself does not seem a plausible explanation for our lack of effect. We would like to note once more that the design of this study was modeled to be as similar as possible to the primate studies. Li & DiCarlo (2008) found substantial alterations of position tolerance after an exposure phase consisting of 400 swap trials and 400 non-swap trials, which is the same number of trials as we used here. We calculated that our experiment had a moderate (0.5986; two-tailed test) to reasonable (0.7238; one-tailed test) power to detect a reduction by half of the size tolerance in LOC, which is the effect size that is to be expected for this amount of exposure based on the studies of Li & DiCarlo (2008; 2010; 2012). Moreover, in the same brain region other anticipated effects – such as effects of size difference, object similarity and object identity – were detected with high significance. Our failure to find an effect of temporal contiguity training thus seems to be due to the absence of the effect rather than a lack of power. At the very least, it suggests that the effect size of changes due to temporal contiguity is much smaller than can be presumed from previous studies.

Apart from being smaller, it might also built up more slowly than expected. We cannot exclude the possibility that an effect might be found using fMRI when the duration of the exposure phase is increased considerably. One observation indicated that the training might have affected the associations between images outside of the visual cortex. In two medial temporal lobe (MTL) regions, namely the hippocampus and parahippocampal gyrus, the neural distance between an object in the medium size and the same object in a different size was smaller in the non-swap condition than in the control condition, implying increased representational similarity because of the temporal association. The neural patterns in these regions were noisy (distances A to D were not significantly larger than zero), and the effect did not survive correction for multiple comparisons. Moreover, no other association effects were found in these regions. Therefore, this observation does not allow us to draw firm conclusions. Future studies in which the imaging parameters are optimized for coverage of the medial temporal lobe, and the hippocampus and parahippocampal gyrus in particular, might bring more clarity.

Another important difference between our study and the primate experiments is the spatial resolution. While the electrophysiological methods used by Li & DiCarlo allowed them to measure responses of single neurons or small groups of neurons and adjust stimulus selection to their selectivity, the smallest measurement unit of fMRI, a voxel, consists of hundreds of thousands of neurons. A computational modeling study by Isik, Leibo, and Poggio (2012) suggested that, depending on the size of the measured cell population and the amount of exposure, large changes at the level of single cells might be negligible at the population level. This could be the case if the effects of the exposure are confined to the small subset of cells with a high preference for one of the objects involved in the training, as was the case in the model of Isik et al. (2012). It is not sure that this can explain the discrepancy with the monkey results, because much of the evidence came from multi-unit recordings that also already pool across multiple neurons. Nevertheless, it is a possible hypothesis that should be investigated further, e.g. by a more unbiased sampling of neurons in a monkey electrophysiology study.

In conclusion, we found that object identity is represented in LOC in a size invariant way. Neural patterns in LOC also contain information about object size and object similarity. We did not find any evidence that temporal contiguity training affects size tolerance in this region. More research focusing on other brain regions, including the medial temporal lobe, incorporating longer training paradigms, and/or investigating how effects are distributed across neuronal populations is needed to further elucidate the role of temporal contiguity in learning invariance.

## Acknowledgements

The data has been made publicly available via the Open Science Framework and can be accessed at https://osf.io/csu34/.

This work was supported by the European Research Council (ERC-2011-StG-284101), a Hercules grant (ZW11_10), the Fund for Scientific Research-Flanders (FWO 11X9918N and G088216N), and the KU Leuven Research Council (C14/16/031).

